# Plant spatial aggregation modulates the interplay between plant competition and pollinator attraction with contrasting outcomes of plant fitness

**DOI:** 10.1101/2022.10.21.513236

**Authors:** María Hurtado, Oscar Godoy, Ignasi Bartomeus

**Affiliations:** Departamento de Biología, Instituto Universitario de Investigación Marina (INMAR), Universidad de Cádiz Puerto Real, E-11510, Spain; Departamento de Ecología Integrativa, Estación Biológica de Doñana, EBD-CSIC, Avda. Américo Vespucio 26, Seville E-41092, Spain

**Keywords:** Neighborhood effect, plant fitness, plant-pollinator interaction, spatial scales, structural equation models

## Abstract

Ecosystem functions such as seed production are the result of a complex interplay between competitive plant-plant interactions and mutualistic pollinator-plant interactions. In this interplay, spatial plant aggregation could work in two different directions: it could increase intra- and interspecific competition, thus reducing seed production; but it could also attract pollinators increasing plant fitness. To shed light on how plant spatial arrangement modulates this balance, we conducted a field study in a Mediterranean annual grassland with three focal plant species with different phenology (*Chamaemelum fuscatum* (early phenology), *Leontodon maroccanus* (middle phenology) and *Pulicaria paludosa* (late phenology)) and a diverse guild of pollinators (flies, bees, beetles, and butterflies). All three species showed spatial aggregation of conspecific individuals. Additionally, we found that the two mechanisms were working simultaneously: crowded neighborhoods reduced individual seed production via plant-plant competition, but they also made individual plants more attractive for some pollinator guilds, increasing visitation rates and plant fitness. The balance between these two forces varied depending on the focal species and the spatial scale considered. Therefore, our results indicate that mutualistic interactions not always effectively compensate for competitive interactions in situations of spatial aggregation of flowering plants, at least in our study system. We highlight the importance of explicitly considering the spatial structure at different spatial scales of multitrophic interactions to better understand individual plant fitness and community dynamics.

## 1. INTRODUCTION

Species fitness, measured as the ability of individuals to contribute with offspring to the next generation, modulates several ecological processes at the community scale such as changes in species relative abundances across years, ultimately defining the maintenance of biodiversity (Hacker & Gaines, 1997; Schmidtke *et al*., 2010). Plant reproductive success is a complex process which is considered to be generally affected by species interactions and environmental conditions. For flowering plants, two key types of biotic interactions are considered. These are competitive interactions due to plant competition for space, nutrients (Tilman, 1990; Craine & Dybzinski, 2013) and shared natural enemies such as herbivores (Hulme,1996) and mutualistic interactions with pollinators which mediate flower’s pollination (Ollerton *et al*., 2011; Thompson, 2006).

Beyond these competitive and mutualistic interactions that affect plant fitness in opposite directions, more subtle effects emerge when we consider explicitly the spatial configuration of plant individuals and their pollinators. For example, the number of floral visitors that a plant receives not only depends on the plant characteristics, but also on the plant neighborhood densities (Ghazoul, 2006; Seifan *et al*., 2014; Bruninga-Socolar & Branam, 2022). Hence, the plant neighborhood can indirectly impact plant reproductive success via pollinator attraction (Lázaro *et al*., 2014; Albor *et al*., 2019; Underwood *et al*., 2020; de Jager *et al*., 2022). Although the outcome of this indirect interactions is hard to predict as it depends on the characteristics of the plant neighborhood (Stoll & Patri, 2001; Underwood *et al*., 2020), the floral preferences of the pollinators involved (Ghazoul, 2006; Hegland & Totland, 2012; Seifan *et al*., 2014; de Jager *et al*., 2022), and their behavior and foraging ranges (Sowig, 1989; Lázaro & Totland, 2010; Seifan *et al*., 2014), we can foresee some contrasting processes.

One the one hand, some species in mixed species neighborhoods can benefit from the effect that particular species, some of them considered magnet species (Thompson, 1978; Seifan *et al*., 2014), have in attracting more pollinators (Carvalheiro *et al*., 2014; Mesgaran *et al*., 2017; Bergamo *et al*., 2020; Bruninga-Socolar & Branam, 2022).

However, these positive spillover effects can turn into competition for pollinators if particular species are less attractive (Mesgaran *et al*., 2017). Indeed, the balance between such positive and negative net effects in mixed neighborhoods is a density dependence process that involves both plant and pollinator abundances. Competition for attracting pollinators can occur either because of high local densities of both conspecific and heterospecific individuals (Ghazoul, 2006; Muñoz & Cavieres, 2008; Dauber *et al*., 2010; Seifan *et al*., 2014), or simply because pollinators are scarce (Lázaro *et al*., 2014).

The characteristics that determine the spatial distribution of the organisms involved in plant-pollinator interactions are multiple. The spatial distribution of plant that determine their density and relative abundance (i.e. the relative abundance of intraspecific versus interspecific neighborhoods) are known to be affected by microclimatic conditions, plant competition and facilitation, dispersal capacity or historical events such as order of arrival (Duflot *et al*., 2014; Gámez-Virués *et al*., 2015). However, pollinators are mobile organisms which may be able to track resources and hence be less constrained in their spatial location (Lander *et al*., 2011; Reverté *et al*., 2019). For example, hover flies are wanderers, but spend more time in resource rich patches (Lander *et al*., 2011), and despite bees being central place foragers, they can track their preferred resource in the landscape (Lázaro & Totland, 2010), sometimes along large distances (López-Uribe *et al*., 2016).

Although we can hypothesize that spatial aggregation of plant-pollinator systems can be modulating plant fitness, a key open question is at which scale it operates (Albor *et al*., 2019; Chase & Leibold, 2002; Underwood *et al*., 2020). Answering whether different processes act at different scales is important to understand how they combine their net effect into plant fitness. For example, plant-plant competition in annual systems is considered to act at small spatial scales (order of centimeters) (Levine & HilleRisLambers, 2009; Lanuza *et al*., 2018). However, plant population dynamics including other processes such as dispersal act at larger scales (order of meters) (Pacala & Silander, 1990; Underwood *et al*., 2020). The scale at which plant community composition modulates pollinator attraction and visitation rates is also multiple. Most pollinators use visual and olfactory cues (Chittka & Thomson, 2001) to select their foraging patches at larger scales, however pollinator functional groups perceive floral resources differently across scales (Albor *et al*., 2019). It has been shown that solitary bees can exploit small flower patches and forage at smaller distances (up to 100 m^2^; Zurbuchen *et al*., 2010; Kendall *et al*., 2022) than social bees (Kendall *et al*., 2022). Conversely, other functional groups such as hoverflies are not such scale dependent (Blaauw & Isaaacs, 2014). In addition, behavior also modifies species foraging patterns at local scales. For example, some pollinators such as bumblebees show floral consistency, meaning that when they land on a specific plant species they visit mostly that species in the patch (Chittka & Thomson, 2001; Lázaro & Totland, 2010) while other groups like muscoid flies or hoverflies are less constant in their visits (Lázaro & Totland, 2010).

Here, we study the effect of spatial aggregation of plant-plant and plant-pollinator interactions on plant fitness (measured as viable seed production) in three annual plant species in a Mediterranean grassland in Doñana National Park (South Spain). Our overall hypothesis is that plant-plant and plant-pollinator interactions change with plant homo- and hetero-specific aggregation levels and affect on opposite ways to plant fitness. While plant competitive effects decrease plant fitness, pollinators increase it. We also hypothesize that the strength of both processes is similar, and therefore, floral visitors can compensate for the negative effect of competition on fitness. Finally, we also hypothesize that these opposing effects occur at different spatial scales. While plant competition occurs at local scales, attraction to floral resources, and therefore an increase in visitation rates occur at larger spatial scales, which is the scale at which most effective pollinators take foraging decisions. These processes at contrasting scales may decouple the positive and negative effects of plant competition and pollinator mutualistic interactions.

## 2. MATERIAL AND METHODS

### 2.1 Study System

We conducted our observational study in Caracoles Estate (2680 ha). This natural system is a salty grassland located within Doñana National Park, southwest of Spain (37°04’01.0”N 6°19’16.2”W). The climate is Mediterranean with mild winters and average 50-year annual rainfall of 550–570 mm with high interannual oscillations. Soils are sodic saline (electric conductivity > 4 dS/m and pH < 8.5) and annual vegetation dominates the grassland with no perennial species present. The study site has a subtle micro topographic gradient (slope 0.16%) enough to create vernal pools at lower parts from winter (November–January) to spring (March–May) while upper parts do not get flooded except in exceptionally wet years (Lanuza *et al*., 2018). Along this gradient (1 km long x 800 m wide), we established in 2015 nine plots, three in the upper part, three in the middle, and three in the lower part. Each plot has a size of 8.5 m x 8.5 m, which is further subdivided in 36 subplots of 1 m^2^ (1 m x 1 m). Average distance between these three locations was 300 m and average distance between plots within each location was 40 m (minimum distance 25 m).

We took advantage of this infrastructure to sample annual plant vegetation and their associated pollinators during 2020. Across plots, we observed 23 co-occurring annual plant species, which represent > 90% of cover. Detailed weekly surveys of pollinators during the flowering season (see below) showed that the flowers of ten of these species were visited by insects, but most of these visits belonging to four different pollinators guilds (bees (14.74%), flies (19.84%), beetles (63.66%), and butterflies (0.8%)) were concentrated (95% of the total of visits) only in three Asteraceae species (*Chamaemelum fuscatum, Leontodon maroccanus* and *Pulicaria paludosa*; Figure A1, APPENDIX A). Therefore, these three species were those considered for further analyses (Table 1). For the analysis butterflies were excluded due to the low visitation to flowers (we only observe 13 visits across species) (Table1).

**Table 1.**
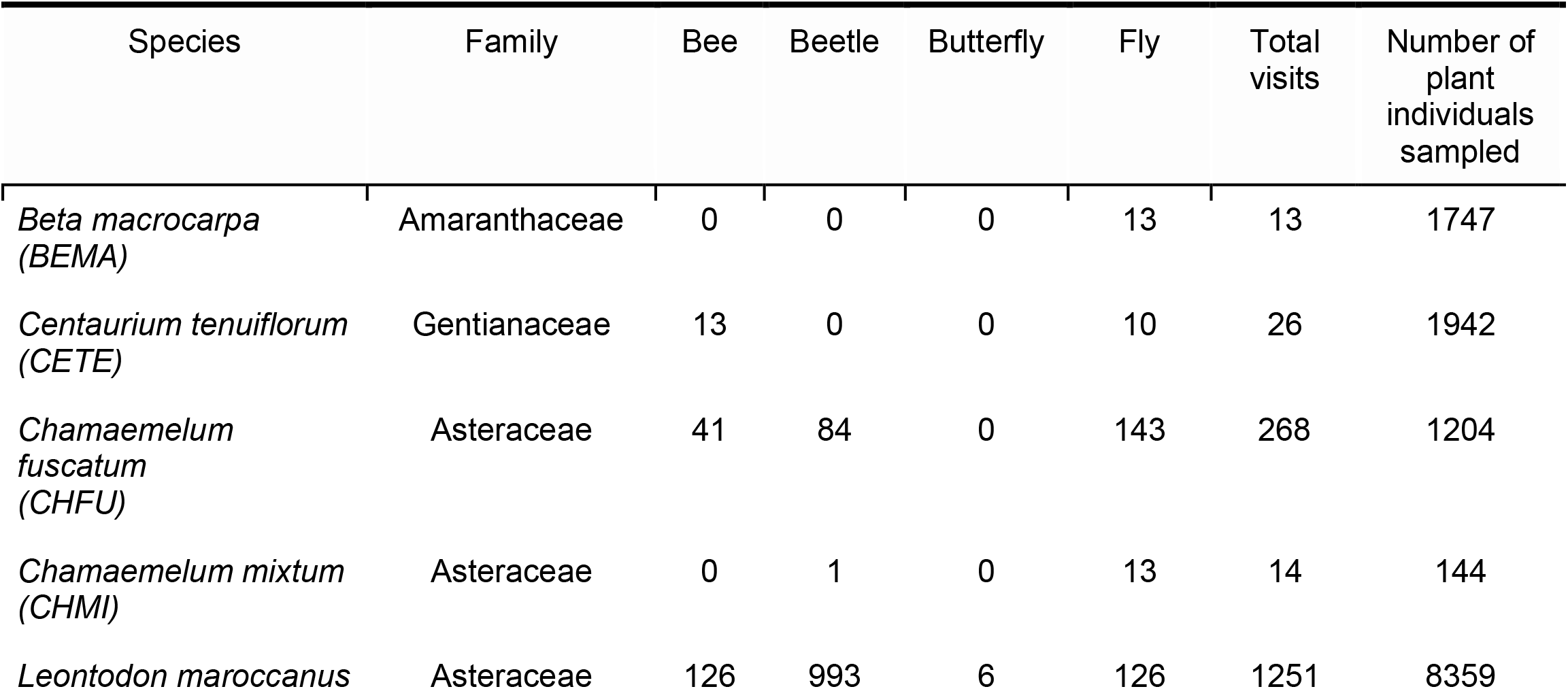
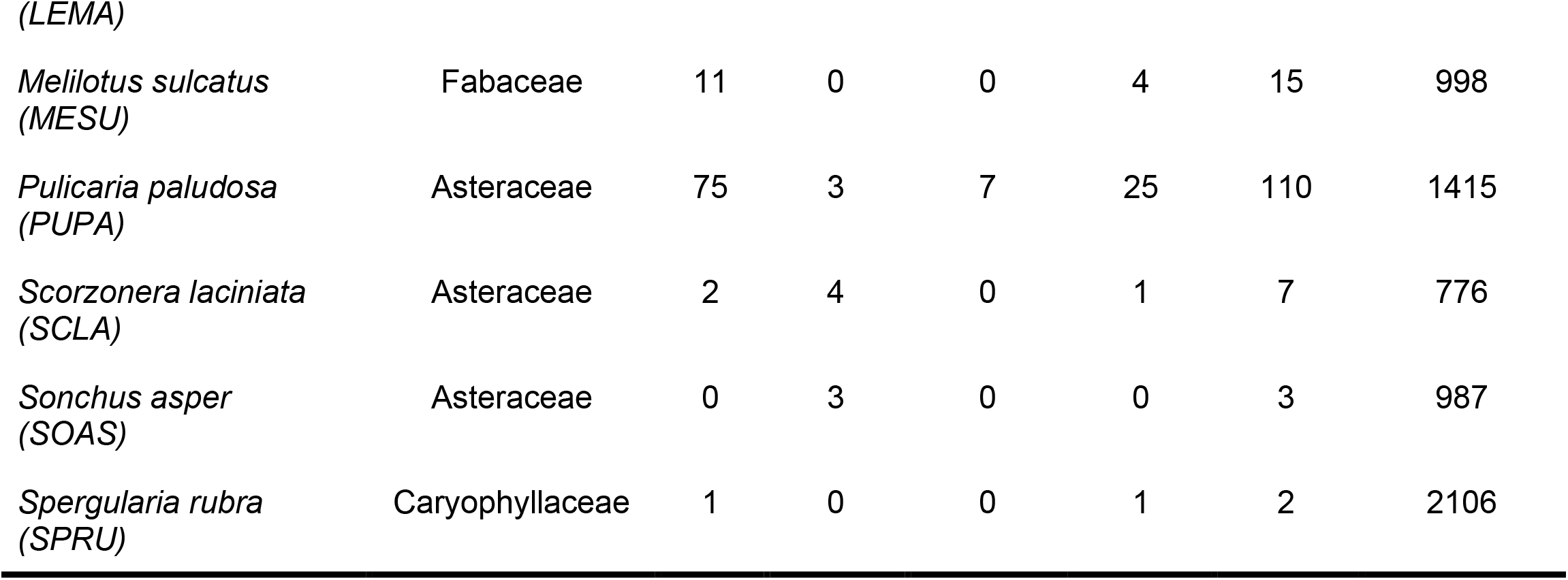
Taxonomic list (and code) in Caracoles field site for those species we observed pollinators visiting during 2020. Specifically, it is shown the number of visits of each pollinator group to each plant species. Note that the abundances of each plant species that we measured at the plot scale (last column) is correlated with their natural abundances in the site study at larger scales. The table of the 23 plant species is in Table A2, APPENDIX A.

| Species | Family | Bee | Beetle | Butterfly | Fly | Total visits | Number of plant individuals sampled |
| --- | --- | --- | --- | --- | --- | --- | --- |
| *Beta macrocarpa*<br>(BEMA) | Amaranthaceae | 0 | 0 | 0 | 13 | 13 | 1747 |
| *Centaureum tenuiflorum*<br>(CETE) | Gentianaceae | 13 | 0 | 0 | 10 | 26 | 1942 |
| *Chamaemelum fuscatum*<br>(CHFU) | Asteraceae | 41 | 84 | 0 | 143 | 268 | 1204 |
| *Chamaemelum mixtum*<br>(CHMI) | Asteraceae | 0 | 1 | 0 | 13 | 14 | 144 |
| *Leontodon maroccanus* | Asteraceae | 126 | 993 | 6 | 126 | 1251 | 8359 |
| (LEMA) |  |  |  |  |  |  |  |
| *Melilotus sulcatus*<br>(MESU) | Fabaceae | 11 | 0 | 0 | 4 | 15 | 998 |
| *Pulicaria paludosa*<br>(PUPA) | Asteraceae | 75 | 3 | 7 | 25 | 110 | 1415 |
| *Scorzonera laciniata*<br>(SCLA) | Asteraceae | 2 | 4 | 0 | 1 | 7 | 776 |
| *Sonchus asper*<br>(SOAS) | Asteraceae | 0 | 3 | 0 | 0 | 3 | 987 |
| *Spergularia rubra*<br>(SPRU) | Caryophyllaceae | 1 | 0 | 0 | 1 | 2 | 2106 |

### 2.2 Pollinator and neighbor composition sampling

Following the spatial explicit design, our overall set of measurements collected involved three main steps. First, we recorded for each observed individual plant, the number of floral visits received by each pollinator guild. Second, we associated these visits with the abundance of plants sampled at different plant scales (neighborhood scale (7.5 cm^2^), subplot scale (1m^2^), and plot scale (3 and 6m^2^)). Finally, to know its reproduction success we measured the number of fruits produced per individual and the viable seed production per fruit.

For the first step, we sampled the number of floral visits and the identity of the guild that each individual plant received. This sampling spanned from the 13th of February to the 18th of July of 2020, which corresponds from the emergence of the earliest flowers of *C. fuscatum* to the latest flowers of *P. paludosa*. Specifically, once per week, we spent 30 minutes per plot, when insect activity is greatest (between 10:00 am and 15:00 am), recording the number of interactions between insects and plants at the subplot level (1m x 1m). To reduce any temporal bias in observations, we randomly select each week which plot was initially sampled. A visit was only considered when an insect touched the reproductive organs of the plants. All pollinators were either identified during the survey or they were net-collected for their posterior identification at the lab. Later, they were grouped into four distinct categories mentioned before: bees, beetles, butterflies and flies (Table A1 in APPENDIX A). Voucher specimens were deposited at Estación Biológica de Doñana (Seville, Spain). Overall, the methodology rendered 54 hours along 19 weeks of sampling. With these field observations, we calculated the total number of visits per pollinator guild in each subplot to each plant species; we assumed that if a pollinator was present in a plot it has the potential to visit all flowering individuals.

For the second step, we measure the number and identity of each plant individual following common procedures of plant competition experiments (Levine & HilleRislambers, 2009; Lanuza *et al*. 2018). Specifically, at the peak of flowering of each species (i.e. when approximately 50% of the flowers per individual were blooming (*C. fuscatum*: early april, *L. maroccanu*s: middle-end April and *P. paludosa* end of May), we chose a focal individual in each subplot for measuring reproductive success, and we used it as the center of a circle with a radius of 7.5 cm, in which the number of individuals and its identity at the species level was recorded. For the three species of our study, we surveyed the neighborhood of 605 individuals. We additionally counted the number of individuals and their identity at the scale of the subplot (1 m^2^) for all species found, which included insect and non-insect pollinated species. Because we measured abundances for each 324 subplot (36 subplots x 9 plots), we were also able to relate to each targeted individual the number of conspecific and heterospecific individuals at larger spatial scales (3m^2^ and 6m^2^ (plot level)). For calculating the neighbors of each focal individual at different scales, we did not consider the subplot edges in order that all focal individuals have the same subplot surrounding them. In total we had the neighbor abundances for each 144 subplots (16 subplots x 9 plots). The survey of abundances across subplots yielded a total of 38220 plant individuals with individual subplots varying between 150 individuals to 1 individual as the minimum, the mean of the individuals that have been counted per subplot is 14 individuals.

In the last step, we sampled for each individual identified at the center of the 7.5 cm^2^ the number of developed fruits and seeds. With this information we measured the reproductive success in two different ways: number of viable seeds per fruit (for now on seed set) and number of fruits per individual (i.e fruit set). The number of fruits per individual was measured in the field as the number of flowers because the three species were Asteraceae. The seed sets were counted at the lab once the fruits were ripped. To account which proportion of the seed set were viable, we visually discarded those that look undeveloped or void. However, measuring the seed set for all fruits of each individual is not feasible for logistic reasons. Therefore, we decided to characterize the species seed set by taking at least one fruit per individual per subplot across the grassland. Such characterization aimed to sample individuals of the three species across the range of floral visits and spatial arrangements observed. In the subplots in which we do not have data for the field (∼59% of the total), we assume that the number of the seed set would be the mean of the seed set of the plot for each species. Note that we observe marked differences in seed set across plots. In total, we sampled across the nine plots 113 fruits of *C. fuscatum*, 199 fruits of *L. maroccanus* and 150 fruits of *P. paludosa*.

### 2.3 Plant pollinator dependance

The net reproductive success of individual plants depends on the number and type of pollinator visits. However, with these field observations, we cannot establish the baseline of which is the reproductive success of our studied species in the absence of floral visitors. Therefore, to assess the degree of self-pollination for each of the Asteraceae species (*C. fuscatum, L. maroccanus* and *P. paludosa*), we conducted a parallel experiment in which we randomly chose twenty floral buttons per species and we excluded pollinators for ten of these covering them by a small cloth bag. For all three plant species, we hypothesize that pollinators could increase their reproductive success, although the rate of increase could vary among species due to selfing processes. The viable and no viable seeds were counted at the lab once the fruits were ripped.

### 2.4 Statistical analysis

To describe the spatial arrangement of pollinators, plant species and their reproductive success we determined the degree of auto spatial correlation by means of Moran’s I test. Briefly, Moran’s I indicate whether the spatial distribution of a response variable across distance is more similar (positive values) or less similar (negative values) than in a random distribution. Moran’s I ranges from -1 to 1, and their associated error (95% confidence interval) is calculated by bootstrapping. Our unit of analysis in the Moran’s I test was the subplot level (all the subplots, 324), and therefore distance among subplots were calculated in meters. For the case of the spatial distribution of plant abundances, we considered the information obtained at 1m^2^, which pooled the sum of counted plant individuals across all 23 species. For individual plant reproductive success, we used the average of the seed set per species across subplots. Finally, for pollinators, we used the abundance of pollinators per guild across subplots (sum of the counts of each floral visitor per subplot).

To evaluate the effect of the spatial arrangement of modulating the opposing effects of plant-plant interaction and plant-pollinator interaction of plant reproductive success, we used Structural Equation Models (SEMs) (Suárez-Mariño *et al*., 2022) with a multigroup analysis context. The multigroup context was used to further test the hypothesis that different processes affect plant reproductive success at different spatial scales. Prior to SEM analysis, we ran Pearson correlations among all predictors to make sure the different analyzed variables were not highly correlated (i.e. r > 0.8). The only variables that are highly correlated are the number of fruits with total viable seed production (0.82; full correlation matrix in Figure A2.A, APPENDIX A). This was an expected result as total viable seed production (i.e total seed set) is the product of the number of fruits multiplied by seed per fruit. Despite this correlation, we kept both predictors because we expected different ecological strategies to maximize reproductive success among species. While some species invest more in flower production at the expense of inverting in individual seeds, other species follow the converse strategy. We also checked the correlation between the different scales at which plant abundance was measured (7.5 cm^2^, 1 m^2^, 3 m^2^ and 6 m^2^), because larger scales have been calculated summarizing the 1 m^2^ scale. We found weak correlations for some neighbor aggregations (Figure A2.B, APPENDIX A), which are important for interpreting the results. Prior to conducting the SEM analysis, we rescaled all the variables to reduce influence of more spread variables.

The causal a priori SEM structure for all our species was the same and considered the following direct and indirect links. First, all pollinator guilds can potentially affect seed reproductive success although the sign can be positive, neutral or negative due to their behavior, while some guilds are truly pollinators such as bees others may be floral and pollen herbivores such as some beetles. Furthermore, we separated the effect of the number of conspecific neighbors on the number of fruits produced (i.e. fruit set) from the effect of overall density (total number of conspecific and heterospecific neighbors). While the former neighborhood type could positively and negatively affect plant reproductive success due to competition or facilitation, the latter neighborhood type would predominantly affect the attraction of floral visitors and therefore the number of visits. We added relations between some exogenous variables (e.g. correlation between different pollinator guilds) as suggested by the model fit (see Eq. (1), Eq. (2) and Eq. (3), APPENDIX A and paths depicted in Figures 2, 3 and 4) when ecologically sensible. In the case of *C. fuscatum* we have added the relation between viable seeds per fruit and heterospecific neighbors, and the correlation between the number of visits of beetles and flies. For *L. maroccanus* we have added the relation between viable seeds with conspecific neighbors, the visits of beetles with fruit set and the correlation between seed set and the total seed set. Lastly, for *P. paludosa* we add the relations between fruit set with fly and bee visits, and the correlations between seed set with the total seed set and the fruit set, and the correlation with fly visits with bee and beetle visits. The addition of these relationships was guided by using the modification index (mi). This index is the chi-squared value, with 1 degree of freedom, by which model fit would improve if we added a particular path or constraint freed. When a mi index is higher than 3.64 means that there is a relation path missing (Whalley, 2019). We assess the goodness of statistical fit for each individual species following by an ANOVA procedure and other relevant indices: root mean squared error of approximation (RMSEA), comparative fix index (CFI), standardized root mean square residual (SRMR) (Kline, 2015).

To test whether the importance of these direct and indirect paths are scale dependent we constructed one model constrained (i.e. all paths are forced to get the same values across scales) and another without constraints (i.e. each path can vary across scales). The spatial scales considered were 7.5 cm^2^, 1 m^2^, 3 m^2^ and 6 m^2^. A constrained model means the intercept of the observed variables and the regression coefficients are fixed across the different scales (i.e. no variation). Within the unconstrained model such variation could occur due to the variation in conspecific, and in the overall number of neighbors across scales. To test which type of model (constrained versus unconstrained) fit best the data, we performed ANOVA and AIC. For *C. fuscatum* (p.value = 0.880; DF= 48; CFI= 1.00; RMSEA= 0.00; SRMR= 0.042) and *L. maroccanus* (p.value= 0.869; DF= 44; CFI= 1.00; RMSEA= 0.00; SRMR= 0.037) the unconstrained model considering a spatial scale effect was more supported (Pr(>Chisq) < 0.001, See Table A3 of the APPENDIX A), while the constrained model better supported *P. paludosa* data (p.value= 0.253; DF= 95; CFI= 0.99; RMSEA= 0.038; SRMR= 0.095). All the p.values of the model selected per each species are not significant (p.value > 0.05) and CFI close to 1, RMSEA < 0.04 and SRMR < 0.1, indicating a good statistical fit (Table A3, APPENDIX A).

Finally, to disentangle the direct effect of plant neighborhoods on total seed set from the indirect effect of plant neighborhoods that is mediated by pollinators visits, we calculated the total, direct and indirect effects by multiplying the coefficients involved in each path. To do this comparison we selected the 7.5 cm scale, as we advance that is the scale at which we observed stronger negative relationships likely due to plant-plant competition. To calculate the direct competitive effects of neighbors we have considered the effect of the intra and inter-neighbors on fruits multiplied by the effect of fruits in the total seed set. To calculate the effect of competition mediated by floral visitors we have considered the effect of the intraspecific and interspecific neighbors on pollinators multiplied by the pollinators effect on seed set and the effect of the seed set on total seed set. In the case where neighbors also affected seed production, these paths were included in the calculation of the direct effects. To calculate the total effects, we have summed the path of competitive effects and the path of the effect mediated by pollinators. Note that estimates in Figures 2, 3 and 4 are rounded, but we used all decimals to calculate direct and indirect paths. The methodology used to calculate the direct and indirect effects are the same used in Bollen (1987) and Grace (2006).

All statistical analyses were conducted with R (R version 4.0.3, 2020-10-10). Moran’s I tests were performed using the packages “spdep” (Bivand & Wong, 2018) and for plotting the results we used the function “moran.plot” for the same package. To rescale the variables we used the “scale” function of R base (Becker *et al*.,1988). Lastly, the structural equation models (SEM) and the multigroup were conducted using the package “lavaan” (Rosseel, 2012) with the “sem” function.

## 3. RESULTS

We observed strong differences and a clear hierarchy in pollinator dependence across our three studied species. *C. fuscatum* was the species that depended most on pollinators, followed by *P. paludosa*, which had a slight dependence and *L. maroccanus* showed no dependence on pollinators. Specifically, the amount of seed set produced by *C. fuscatum* increases by 64% under the open pollination treatment compared to the bagged flowers (mean difference among treatments (Effect size) = -64.07; p-value < 0.002). *P. paludosa* showed not significant changes under open pollination (Effect size= -3.24; p-value= 0.56) yet the number of total seeds is very low in both cases (without pollinators= 49.88 ± 31.32 (mean ± sd); with pollinators= 34.7 ± 14.29) comparing with the other species (Figure A3, APPENDIX A), potentially indicating that pollination could be insufficient in the study area, rather than selfing mechanisms. Finally, *L. maroccanus* produces a large number of seeds in both the pollinator exclusion treatment and the open pollination treatment (Effect size= -8.30; p.value= 0.63), indicating no pollinator dependence (Figure A3, APPENDIX A).

The three species (Figure 1) showed a significant degree of spatial autocorrelation (Moran’s I = ∼ 0.4; p.value= 0.01). Generally, they are fairly aggregated at small distances, but this aggregation decays after the first 50 or 100 meters. Nonetheless, the degree of spatial aggregation of floral visitors, despite significant, was much smaller than that of the plant species (Motan’s I < 0.35; p.value= 0.01; Figure 1), especially for mobile organisms such as flies (Moran’s I = 0.19) and bees (Moran’s I = 0.07; p.value= 0.01; Figure 1). The reproductive success of individual plants showed a similar spatial autocorrelation for the three species than the plant individuals (Moran’s I = ∼ 0.3; p.value= 0.01; Figure A4, APPENDIX A). This means that the reproductive success for the plants is unequal in relation to their spatial distribution.

**Figure 1.**
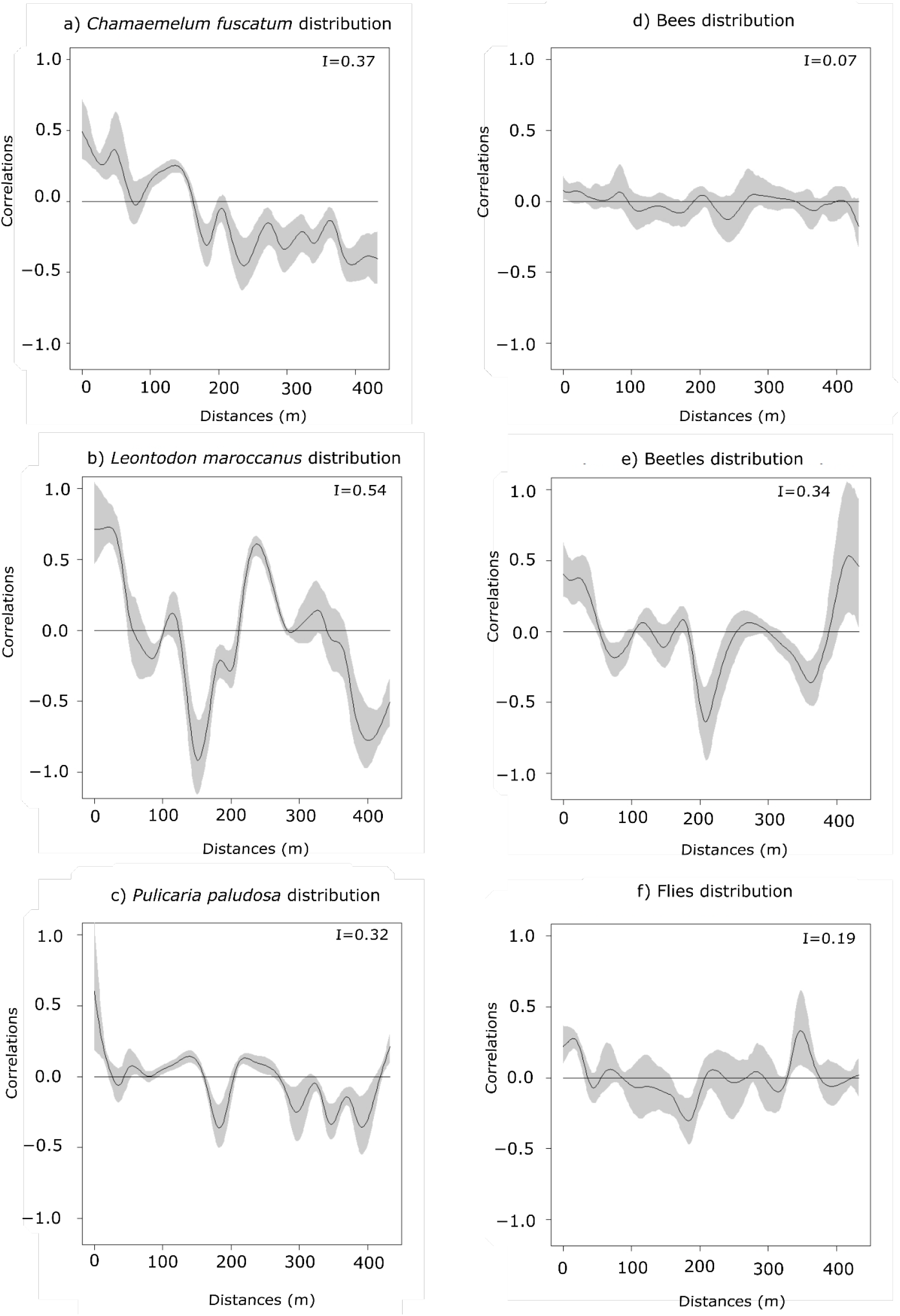
Spatial autocorrelation of plant abundances of the three main species (plots a, b and c: *C. fuscatum, L. maroccanus* and *P. paludosa*, respectively), and the three main pollinators (plots d, e and f: bees, beetles and flies, respectively) at increasing distances. The black line is the spatial correlation value that a species has for each distance, the grey shadow indicates the 95% of the confidence interval. The distribution of plant species individuals is more heterogeneous than the pollinators distribution. The I values are the result of the Moran’s I statistic.

The most important findings when comparing results from the Structural Equation Models (SEMs) is that the reproductive success of the three plant species depended on a different combination of direct and indirect paths, which indicates that there is variability in the biological strategies followed by each species. The best fitted structure of the path diagram revealed that the total number of fruits have a larger influence on the total seed production than the seed set, except in the case of *P. paludosa*. Comparing the direct interactions between plant neighbors (conspecific and heterospecific) and total seed set for *C. fuscatum* and *L. maroccanus* we found a negative relation between the density of conspecific neighbors and fruit production (Figures 2 and 3). Moreover, the effect of conspecific neighbors on the fruit set produced per individual varies depending on the scale. For both species, we can see that the effect of conspecific neighbors on fruits switch across scales. While for *C. fuscatum* is positive at small scales in *L. maroccanus* switches from negative to positive at larger distances. Finally for *P. paludosa*, the effect of conspecific neighbors on the fruit set is negative while the effect of heterospecific neighbors is positive but weak (Figure 4). The neighbors (both conspecific and heterospecific) effect in seed set (in most cases indirect effect through pollinators) and in fruit set is variable depending on the species, in the case of *L. maroccanus* there is a stronger effect of the conspecific neighbors on reproductive success due to its neighbors also affects the seed set, and in the case of *C. fuscatum* the stronger effect is due to the heterospecific neighbors. The role of pollinators in these plant species is in general weak, except in the case of *P. paludosa*, where bees have an important effect on plant reproduction success. However, the number of fruits per plant in the case of *L. maroccanus* and *P. pulicaria* have an effect also in the attraction of pollinators. More fruits (i.e. more flowers per individual), attract more visits of certain pollinators.

**Figure 2.**
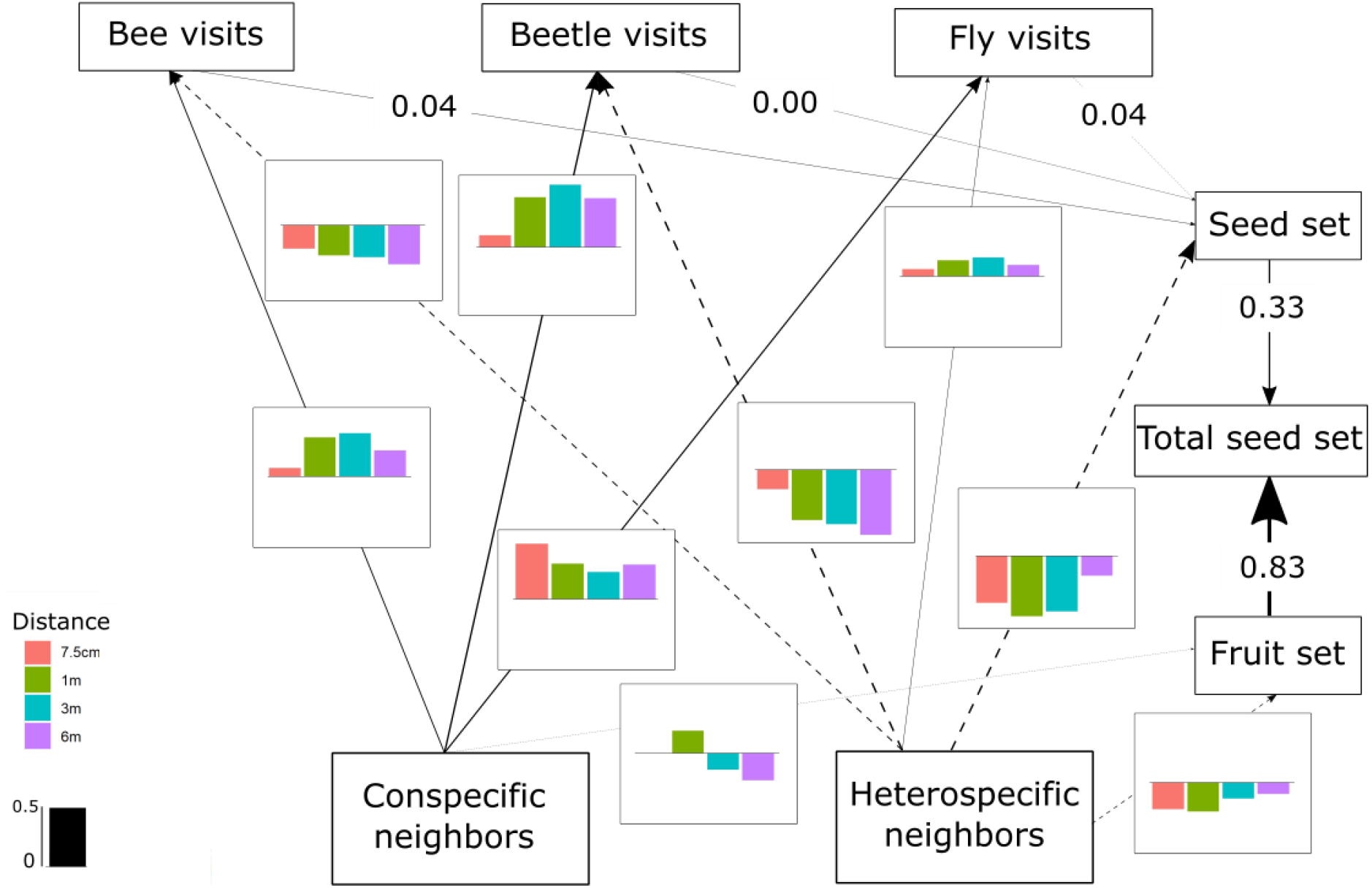
The SEM of *C. fuscatum* which includes the differences in the interactions between scales. Seed refers to the seed set, fruit refers to the number of fruits and total seeds is the total seed set. The lines (dashed and full lines) are proportional to the magnitude of the relation (when different scales, we plot the mean of the standardized total effects across scales) to exemplify the path. The dashed lines are the negative relations. The numbers are the standardized total effects in those variables that remain constant across scales. These barplots show all the standardized total effects of each relation of the model across the different scales. If the value of the barplot is positive, it means that it has a positive effect and if it is negative means that it is a negative effect. It is important to mention that the correlations between the variables are not visualized in the path, but in the SEM model they are included (Eq. (1), APPENDIX A) (p.value = 0.880; DF= 48; R^2^ of total seed set= ∼0.82).

**Figure 3.**
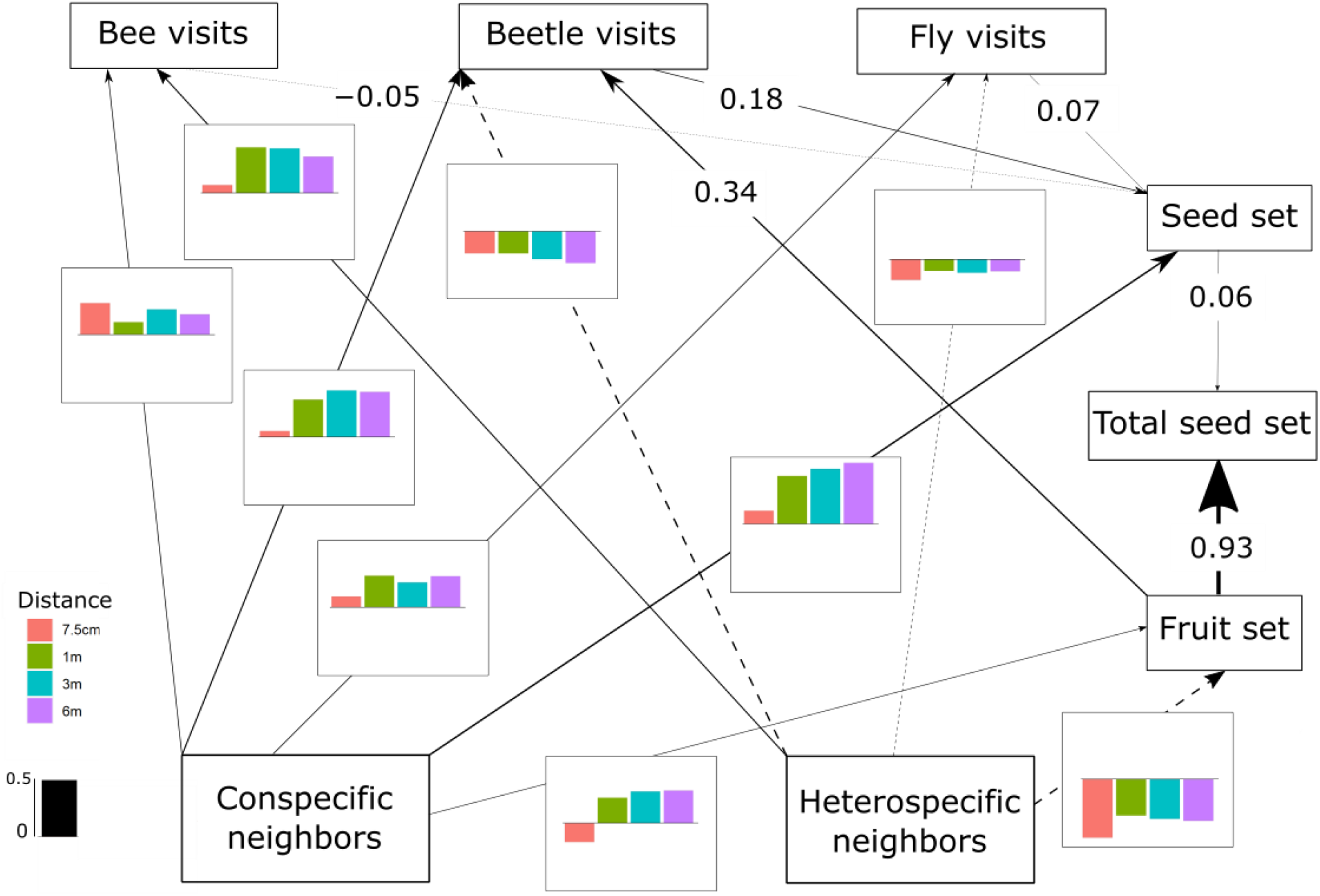
The SEM of *L. maroccanus* which includes the differences in the interactions between scales. The lines (dashed and full lines) are proportional to the magnitude of the relation (when different scales, we plot the mean of the standardized total effects across scales) to exemplify the path. The dashed lines are the negative relations. The numbers are the standardized total effects in those variables that remain constant across scales. These barplots show all the standardized total effects of each relation of the model across the different scales. If the value of the barplot is positive, it means that it has a positive effect and if it is negative means that it is a negative effect. It is important to mention that the correlations between the variables are not visualized in the path, but in the SEM model they are included (Eq. (2), APPENDIX A) (p.value= 0.869; DF=44; R^2^ of total seed set = ∼ 0.91).

**Figure 4.**
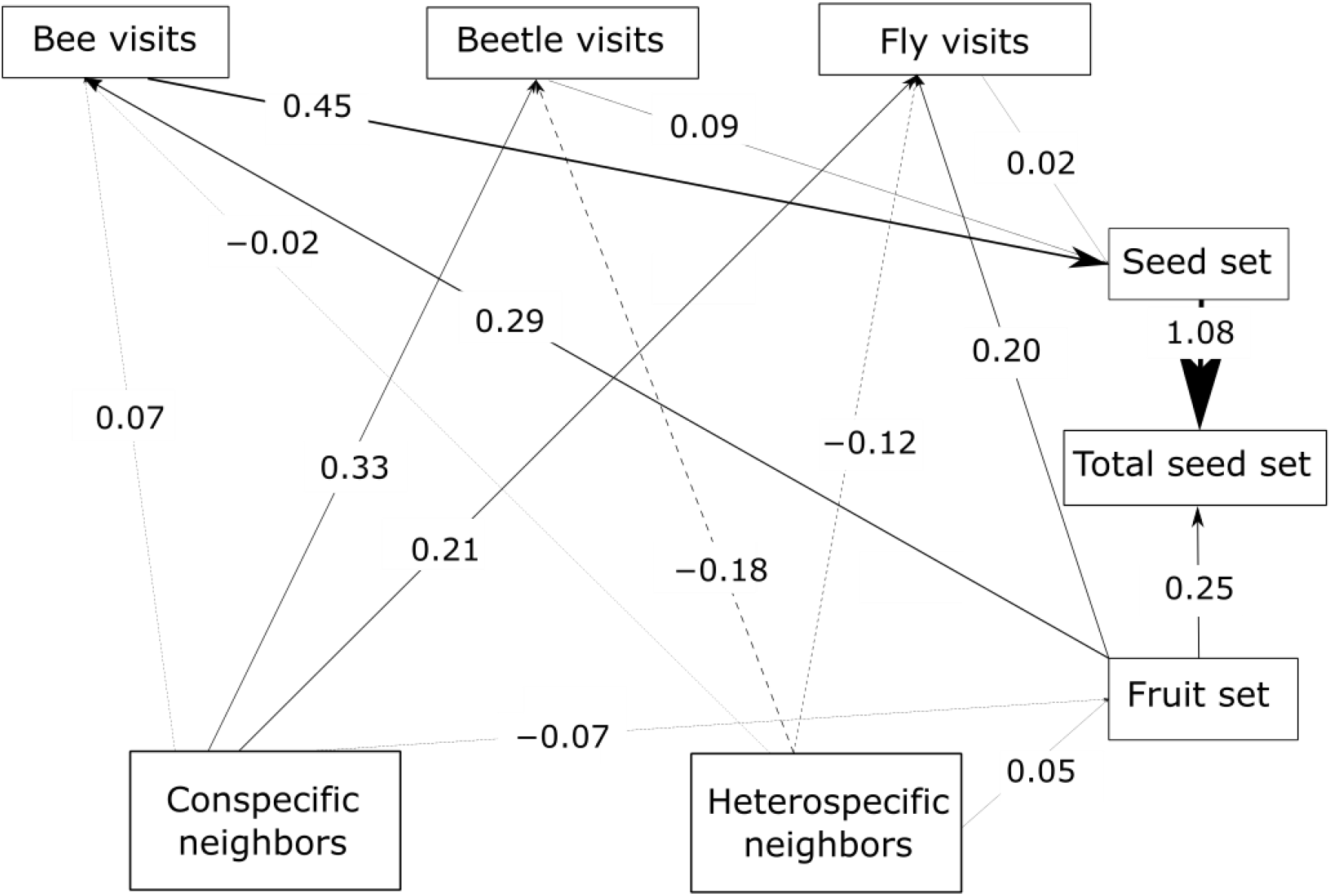
The SEM of *P. paludosa* which includes the differences in the interactions between scales. The lines (dashed and full lines) are proportional to the magnitude of the relation (we plot the standardized total effects) to exemplify the path. The dashed lines are the negative relations. The numbers are the standardized total effects. It is important to mention that the correlations between the variables are not visualized in the path, but in the SEM model they are included (Eq. (3), APPENDIX A) (p.value= 0.253; DF= 95; R^2^ of total seed set= ∼0.4).

We also found a clear effect of the number of both conspecific and heterospecific neighbors on attracting pollinators. Generally, the conspecific neighbors benefit the focal species by attracting more pollinators at medium and large scales, but the effect of heterospecific neighbors is more variable. While heterospecific neighbors always affect the beetle visits negatively, they positively affect the bees in *L. maroccanus* and flies in *C. fuscatum*, but in *P. paludosa* there is a negative effect on the three pollinator groups. When we look at the mean effects of the competition and pollinator mediated paths (Table 3; see effect decomposition across scales in Table A4 APPENDIX A for *C. fuscatum* and *L. maroccanus*) we observed that the positive effect of increased pollinator attraction only compensates for the negative effect of plant competition in *P. paludosa*.

**Table 3.** The direct effects (standardized total effects) of plant competition and the indirect effects mediated by pollinators into the plant reproductive success at the scale of 7.5 cm^2^ (See Table A4, APPENDIX A for the effects on each scale). We have chosen this scale because it is the scale more representative for the path.

| Species | Total effect | Competition effect | Pollinators effect |
| --- | --- | --- | --- |
| *C. fuscatum* | -0.217 | -0.227 | 0.010 |
| *L. maroccanus* | -0.588 | -0.582 | -0.006 |
| *P. paludosa* | 0.023 | -0.003 | 0.027 |

## 4. DISCUSSION

Our most important finding is that the spatial context affects how plant-plant interactions and plant-pollinators interactions contribute to plant reproductive success. Following our main hypotheses, we observed that plants were more aggregated in space than its floral visitors, and they affected in opposite ways plant reproduction success. While plant neighborhoods have a negative effect on plant reproductive success, pollinators result in a more variable, but overall positive effect. However, when comparing the net effect of both sources of plant reproduction success, interestingly we found the positive effect of pollinator visits mediated by the attraction of plant neighbors at larger scales did not compensate for the direct negative effect at neighborhood scales of plant competition in two out of the three studied plants.

Following prior theoretical and observational work, we observed that plant densities, and particularly those of conspecific individuals, had the strongest negative effect on plant reproductive success through a strong effect on fruit set. We interpret this negative effect as competition for common resources such as water, nutrients, or light as well as shared natural enemies (Underwood *et al*., 2020), yet, we acknowledge that we did not explore the ultimate sources of the observed competition. Another important finding is that the scale at which competition acts was different from which the scale pollinators were attracted. Namely, our results suggest that competition effects are stronger at lower scales (Antonovics & Levin, 1980), and confirm that measuring neighborhoods at 7.5 cm^2^ captures the strongest signal of competition (Levine & HilleRisLambers, 2009; Mayfield & Stouffer, 2017; Lanuza *et al*., 2018). However, distances at which pollinators are attracted remains less understood. In our case, pollinator attraction and its further positive contribution on plant reproductive success through pollination visits occur at larger scales up to 3 m^2^.

Indeed, the scale at which different ecological interactions are relevant might differ in other systems. Our study shows that this is a complex interplay between the intrinsic ability of plants to produce seeds in the absence of pollinators, to produce flowers, and therefore to attract pollinators, and the pollinator behavior and their pollination efficacy. In our study, this is exemplified by the contrasted strategies we observed among the three studied species. For instance, *L. maroccanus* and *C. fuscatum* were not limited in the contribution of pollinators to plant reproductive success because *L. maroccanus* is highly self-compatible, and *C. fuscatum* showed no pollen limitation because relied on a high number of visits by small flies which ensure a large seed set across the area. In contrast, the pollination of *P. paludosa was* limited by the low number of bee visits that contributed significantly to increase its reproductive success. This small number of visits could be due to the fact that *P. paludosa* is a late flowering phenology species whose phenology mismatches with the phenology of bees, the fact that *P. paludosa* is not a strongly aggregated species that could attract bees by itself, or maybe it could be simply because bees are scarce in our system. Regardless of these different possibilities, our study shows that the effect of pollinators on plant reproductive success is a spatial explicit process which in turn interacts with the plant and pollinator biology, and despite it might contribute to plant reproductive success positively, it cannot be enough to compensate the negative effects on plant competition in spatially structured environments.

For all species, both plants and pollinator guilds we observed a significant pattern of spatial aggregation, although the magnitude greatly varied across species. Spatial aggregation of plant species is considered to be mediated by a combination of local dispersal and strong preferences for certain environmental conditions (e.g water availability) (Stoll & Patri, 2001). Many annual Asteraceae plant species such as *C. fuscatum* and *P. paludosa* neither possess particular dispersal structures (e.g. pappu) (Howe & Smallwood, 1982; Venable & Levin, 1983) nor are attractive and big enough to be dispersed by seed disperses such as insects or ants (Handel & Beattie, 1990; Rogers *et al*., 2021), therefore they tend to fall in the ground close to their mothers (Venable & Levin, 1983). Other species with pappus structures, such as *L. maroccanus* in this study, can be wind or water dispersed over long distances across space, and their strong spatial aggregation can be due to the selection of particular microenvironmental conditions (e.g substrate) that allow seed germination and establishment (Venable & Levin, 1983; Nathan & Muller-Landau, 2000). For floral visitor guilds, wild bees are known to be central place foragers, which forage close to their nest (Gathman & Tscharnte, 2002) while flies instead seems to have an unspecialized pattern in which they forage distinct flowers along long distances (Inouye *et al*.,2015). Beetles tend to visit less flowers and to stay more time per each flower than the other guilds, having a more clustered aggregation (Primack & Silander, 1975). These arrays of mechanisms suggest that in general it is more likely to find spatial aggregation in plants than in floral visitors. Yet, for any procedure the spatial aggregation is broken, then the remaining question is whether the hierarchy we observed of negative competition effects being stronger than positive mutualistic effects still holds. Future research could manipulate the spatial aggregation across scales to mechanistically test the relative importance of both plant-plant and plant-pollinator interactions for plant reproductive success in spatial uncorrelated environments.

Together, our study provides clear evidence that spatial aggregation across scales, from very small neighborhoods to plot scales is key to determining the magnitude of multitrophic interactions modulating plant reproductive success. Such correlation in conspecific individuals across scales connects pollinator attraction and therefore the mutualistic effect of floral visits (Ghazoul, 2006; Bruninga-Socolar & Branam, 2022; de Jager *et al*., 2022) with the negative competitive effect of dense local neighborhoods (Albor *et al*., 2019; Underwood *et al*., 2020). This connection highlights the fact that the fate on individual reproductive success and therefore the persistence of populations is not only a matter of the degree of temporal autocorrelation (e.g. Lyberger *et al*., 2021; Martinović et *al*., 2021) but also the degree of spatial autocorrelation. However, the spatial effects here documented are rare, and therefore, we call for a need to better integrate observational data with solid theory that connect plant-pollinator systems with multiple trophic interactions in a more comprehensive framework of plant population dynamics. Such integration is paramount because in our study we highlight that predicting the net effect plant-plant and plant-pollinator interactions on plant reproductive success in spatially structured environments is complex, as it results from the combination of pollinators (Underwood *et al*., 2020) and plant characteristics (de Jager *et al*., 2022). We conclude that a more realistic understanding of the direct and indirect effects by which pollinators contribute to plant fitness need to explicitly consider the spatial structure in which these interactions occur.

## DATA AVAILABILITY

The data used to generate the results of this study is deposited at Zenodo https://zenodo.org/record/7216774#.Y07lC3bMK3A

## AUTHOR CONTRIBUTION

OG and IB design the study. MH, OG, IB conducted fieldwork. All authors analyzed the results, and MH and IB wrote the manuscript with substantial contributions from OG.

## ACKNOWLEDGEMENTS

M.H acknowledges financial support provided by the Spanish Ministry of Science and Innovation through the FPI grant (PRE 2019-088280). O.G acknowledges financial support provided by the Spanish Ministry of Science and Innovation through Ramón y Cajal program (RYC 2017-23666) and Proyectos de generación de conocimiento (PID2021-127607OB-100).

## This is the APPENDIX A

**Table A1.** Floral visitor frequency. This is the list of the most accurate identification (ID) of the floral visitors that we have made. Each ID has associated the number of visits in total that we recorded in the field. We classified the ID in four groups of floral visitors: Bee, Beetle, Butterfly and Fly.

| Group | ID of the specimens | Number of total visits |
| --- | --- | --- |
| Bee | *Andrena argentata* | 4 |
| Bee | *Andrena humilis* | 76 |
| Bee | *Andrena* sp | 56 |
| Bee | *Eucera* sp | 4 |
| Bee | *Lasioglossum immunitum* | 2 |
| Bee | *Lasioglossum malachurum* | 104 |
| Bee | *Lasioglossum* sp | 9 |
| Bee | *Osmia ligurica* | 14 |
| Beetle | Family Anthicidae | 9 |
| Beetle | *Brassicogethes* sp | 701 |
| Beetle | *Cassida* sp | 1 |
| Beetle | Family Cerambycidae | 2 |
| Beetle | *Cryptocephalus* sp | 5 |
| Beetle | Family Curculionidae | 1 |
| Beetle | Family Elateridae | 10 |
| Beetle | *Lagorina sericea* | 3 |
| Beetle | *Malachius bipustulatus* | 9 |
| Beetle | Melyridae | 1 |
| Beetle | Mordellidae | 16 |
| Beetle | Oedemeridae | 12 |
| Beetle | *Phaedon* sp | 1 |
| Beetle | *Psilothrix viridicoerulea* | 317 |
| Butterfly | *Euchloe crameri* | 1 |
| Butterfly | Geometridae | 2 |
| Butterfly | *Lasiocampa trifolii* | 5 |
| Butterfly | *Pieris brassicae* | 3 |
| Butterfly | *Vanessa cardui* | 2 |
| Fly | *Anastoechus* sp | 44 |
| Fly | *Bombilus major* | 13 |
| Fly | Family Calliphoridae | 6 |
| Fly | *Cylindromyia* sp | 9 |
| Fly | *Dilophus* sp | 4 |
| Fly | Genus Diptera | 1 |
| Fly | *Empis* sp | 2 |
| Fly | *Episyrphus balteatus* | 14 |
| Fly | *Eristalis* sp | 3 |
| Fly | *Eupeodes corollae* | 1 |
| Fly | *Lomatia* sp | 9 |
| Fly | *Musca* sp | 44 |
| Fly | *Nemotelus* sp | 6 |
| Fly | *Sarcophaga* sp | 23 |
| Fly | *Sphaerophoria scripta* | 32 |
| Fly | Family Syrphidae | 3 |
| Fly | Family Ulidiidae | 122 |

**Figure A1.**
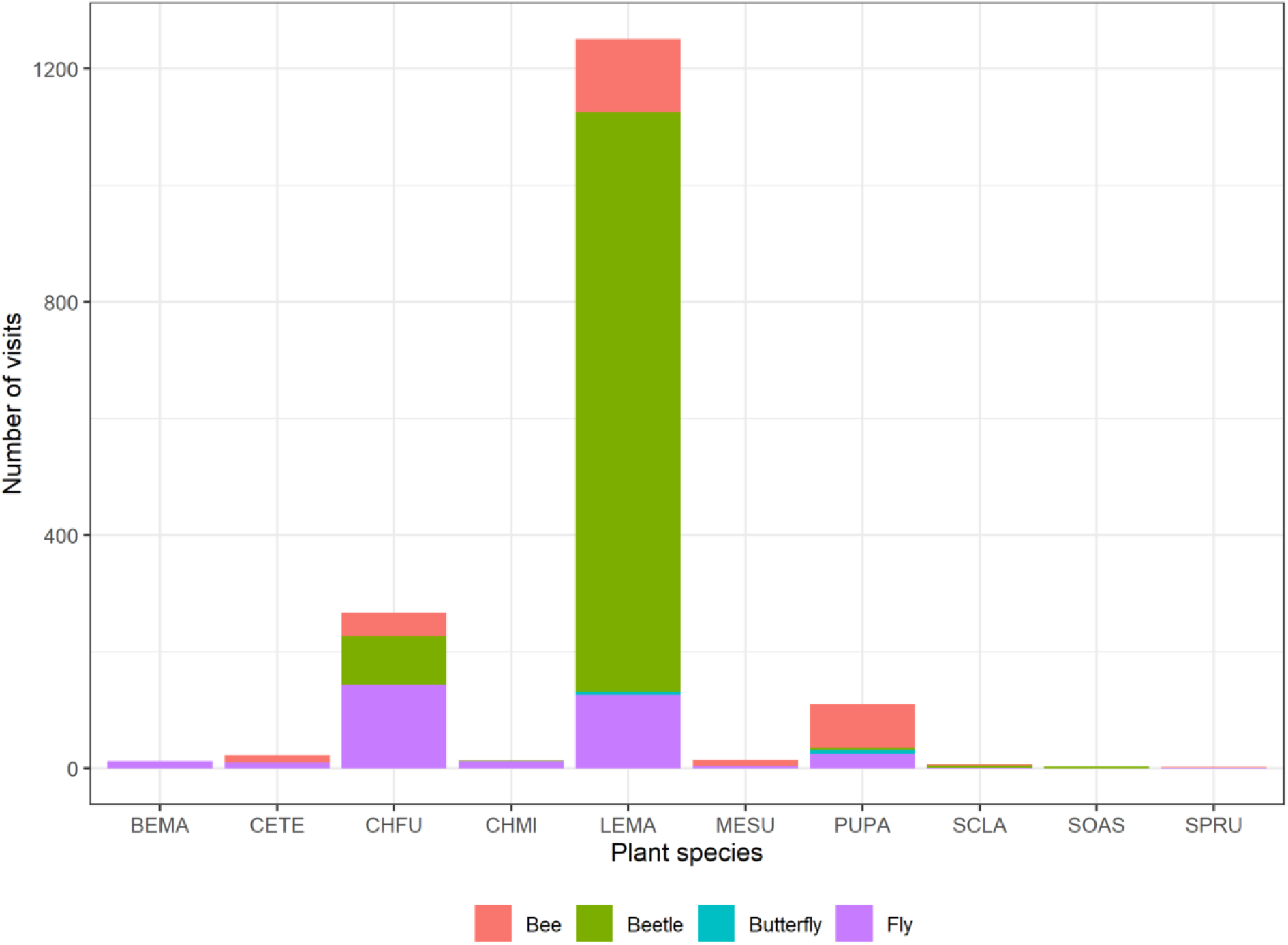
This boxplot shows how the floral visitors are distributed across the plant species. We can observe that the most visited species are *C*.*fuscatum, L*.*maroccanus* and *P*.*paludosa. C*.*fuscatum* is visited mostly by flies, *L*.*maroccanus* is visited mostly by beetles and lastly, *P*.*paludosa* is visited mostly by bees.

**Table A2.**
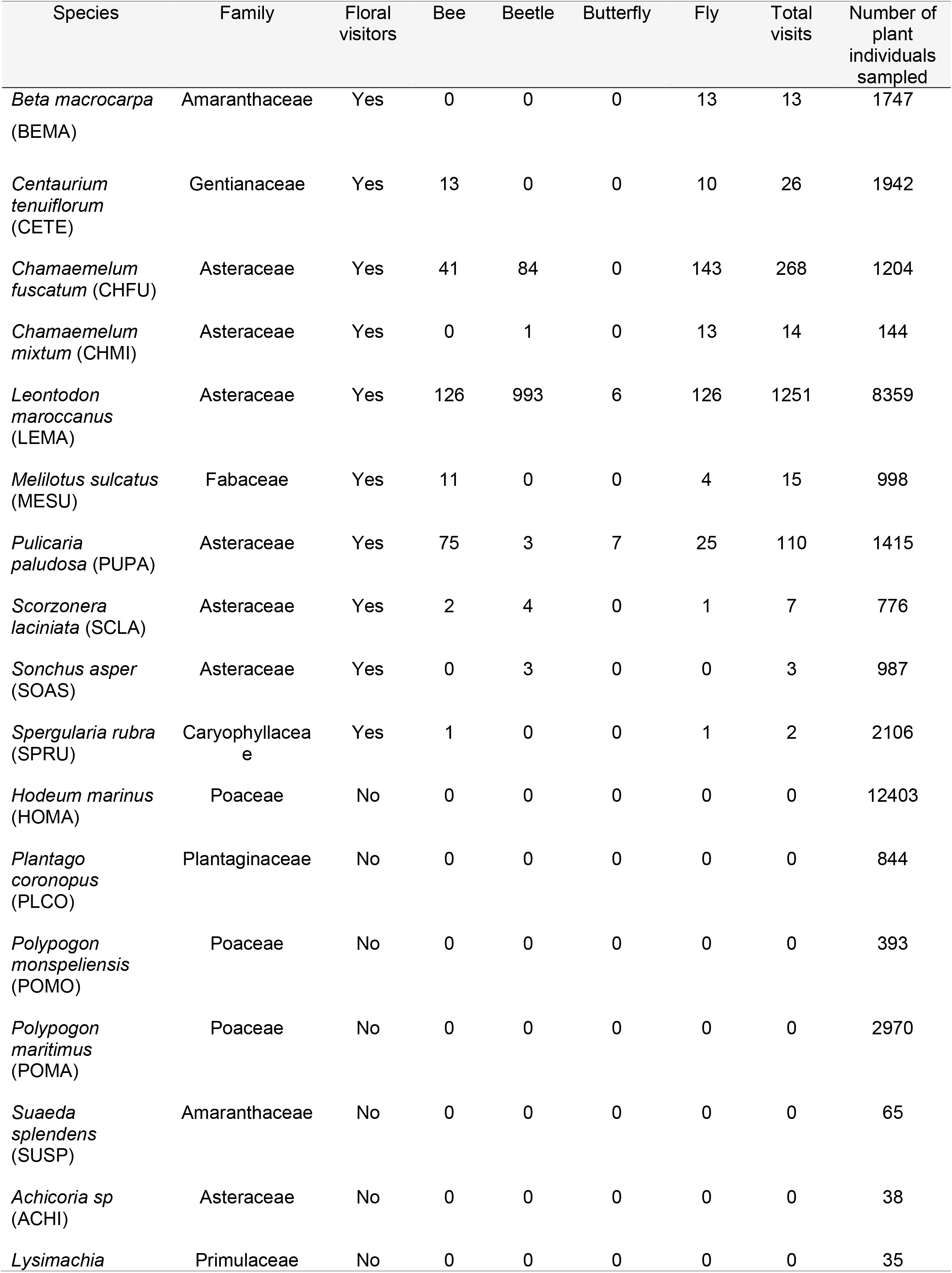
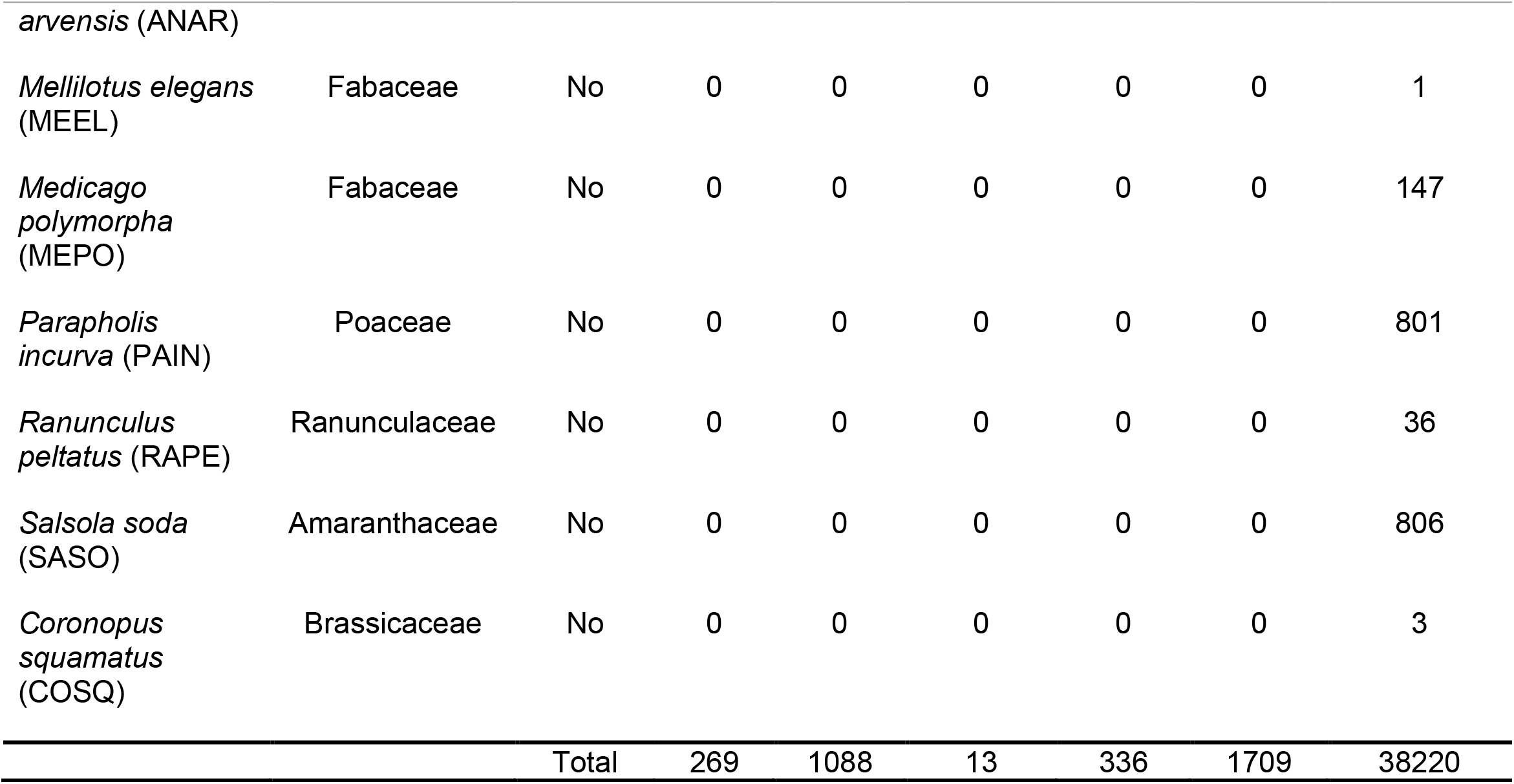
List of species observed in Caracoles Estate in 2020. Code and taxonomic information of the plant species is provided. Also, it is recorded the number of visits of each floral visitor group that receives each plant species. Sample sizes represent the abundances of each species that we measured in the field, and it is correlated with their natural abundances in the site study. In this data the butterflies visits are included, however, due to the low number of visits of that group (only 13 visits) we decided to exclude this data for further analysis.

**Figure A2.**
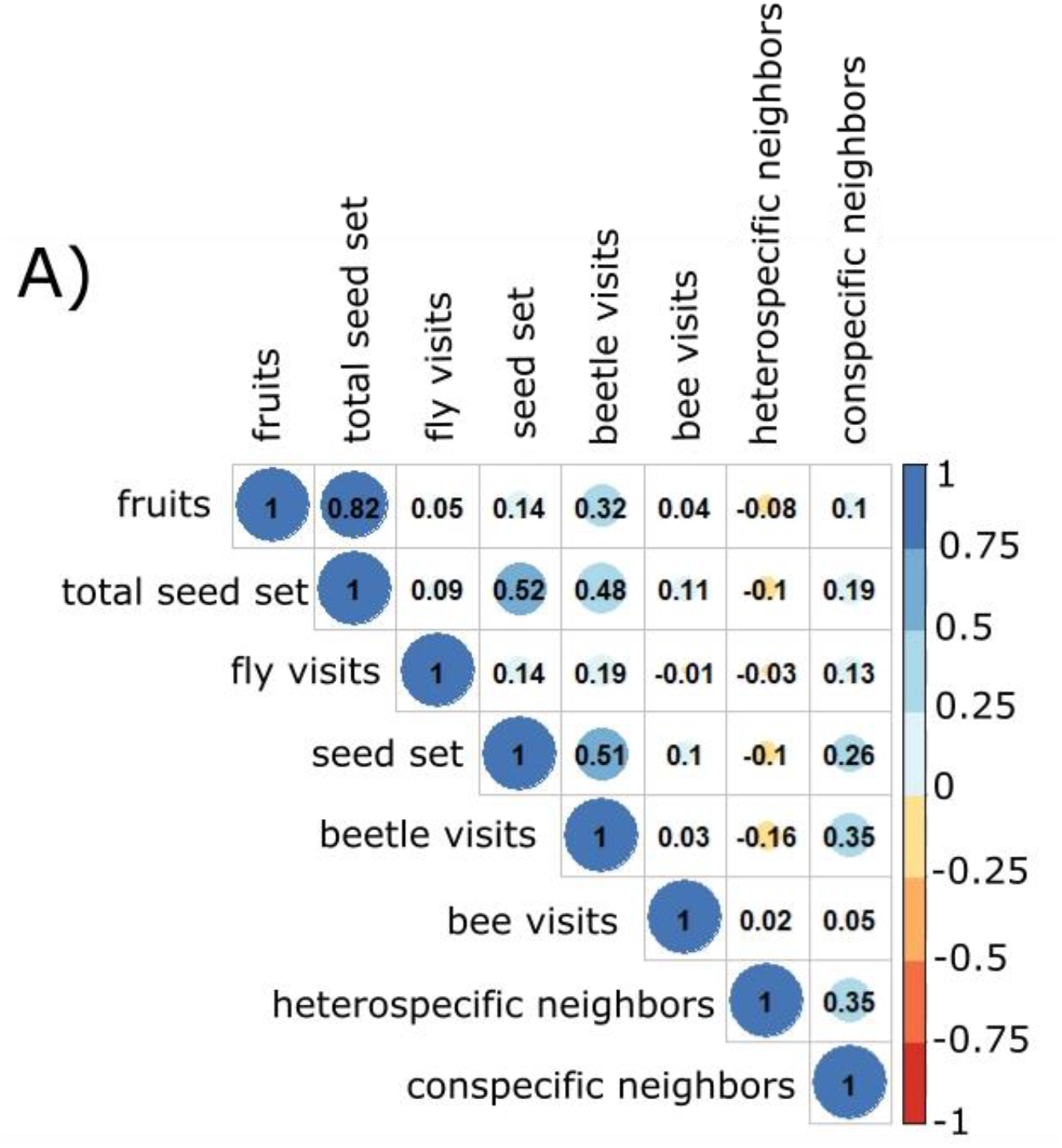
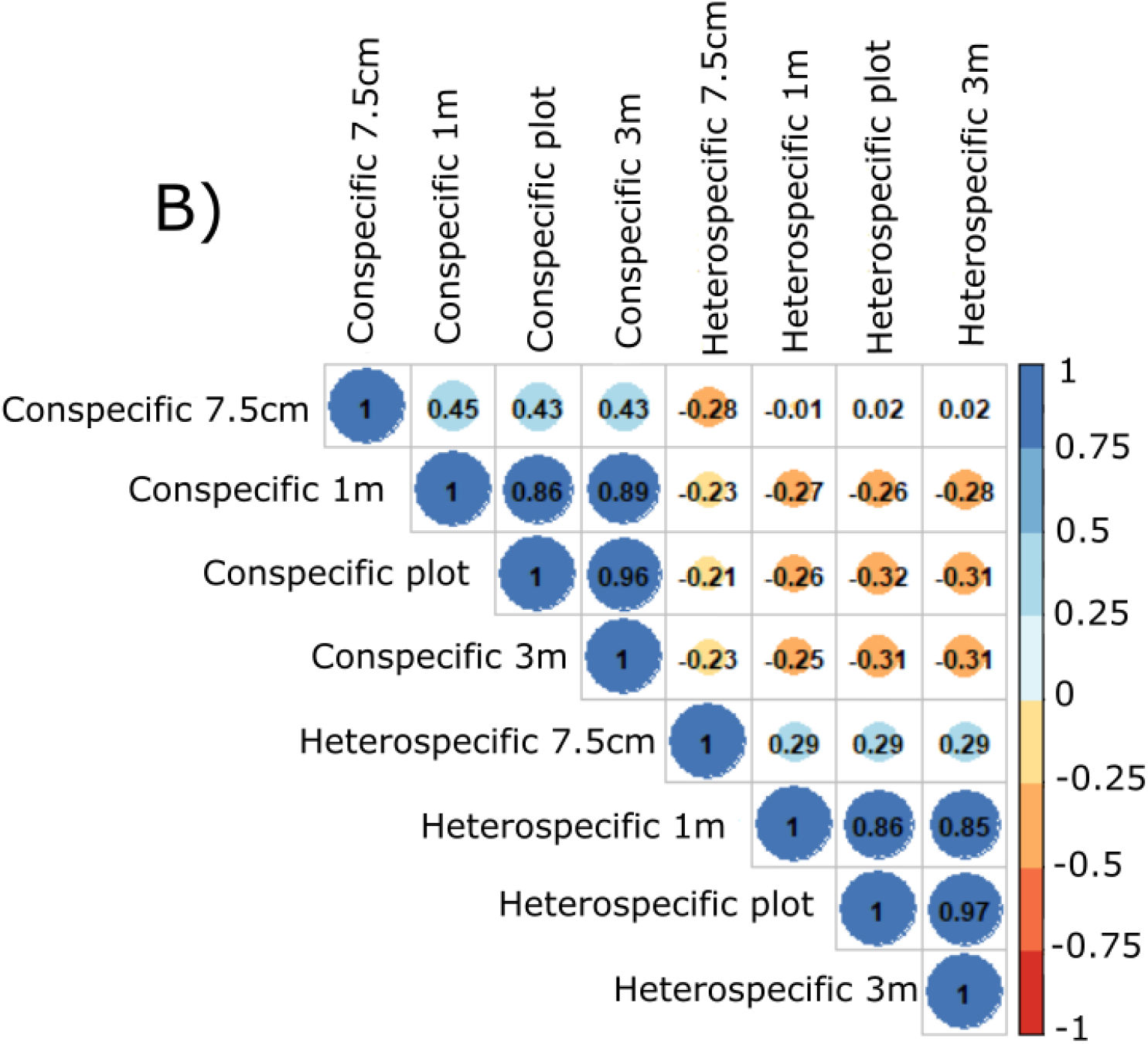
These plots show the correlations between the different variables. In plot A there are the correlations between all the variables included in the model per the three species and in plot B there are the correlations between the different scales of neighbors (7.5 cm^2^, 1m^2^, 3m^2^ and 6m^2^ (plot level)). The strong colors of the cells indicate that there is a strong correlation, and the light colors mean the opposite, there is a slight correlation.

**Figure A3.**
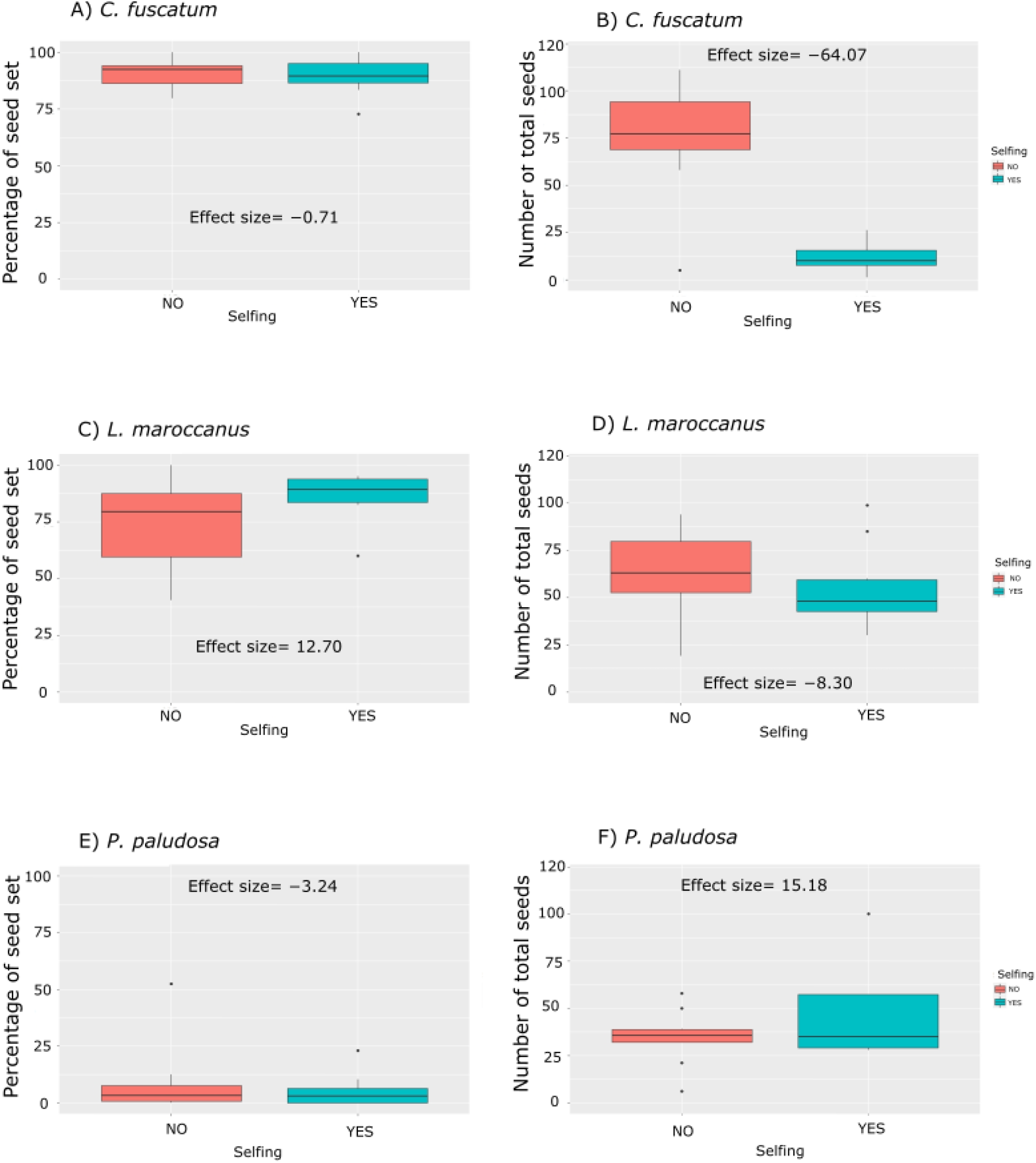
This figure shows the different boxplot for each plant species considering the seed set and the total seed set of the sefing experiment (with or without pollination). In the first column of the plots, we have the percentage of total seed set per species per treatment, and in the second column we have the number of total seeds (viable and no viable seeds) per species and per treatment. The numbers that appear inside the plot are the Effect sizes.

**Figure A4.**
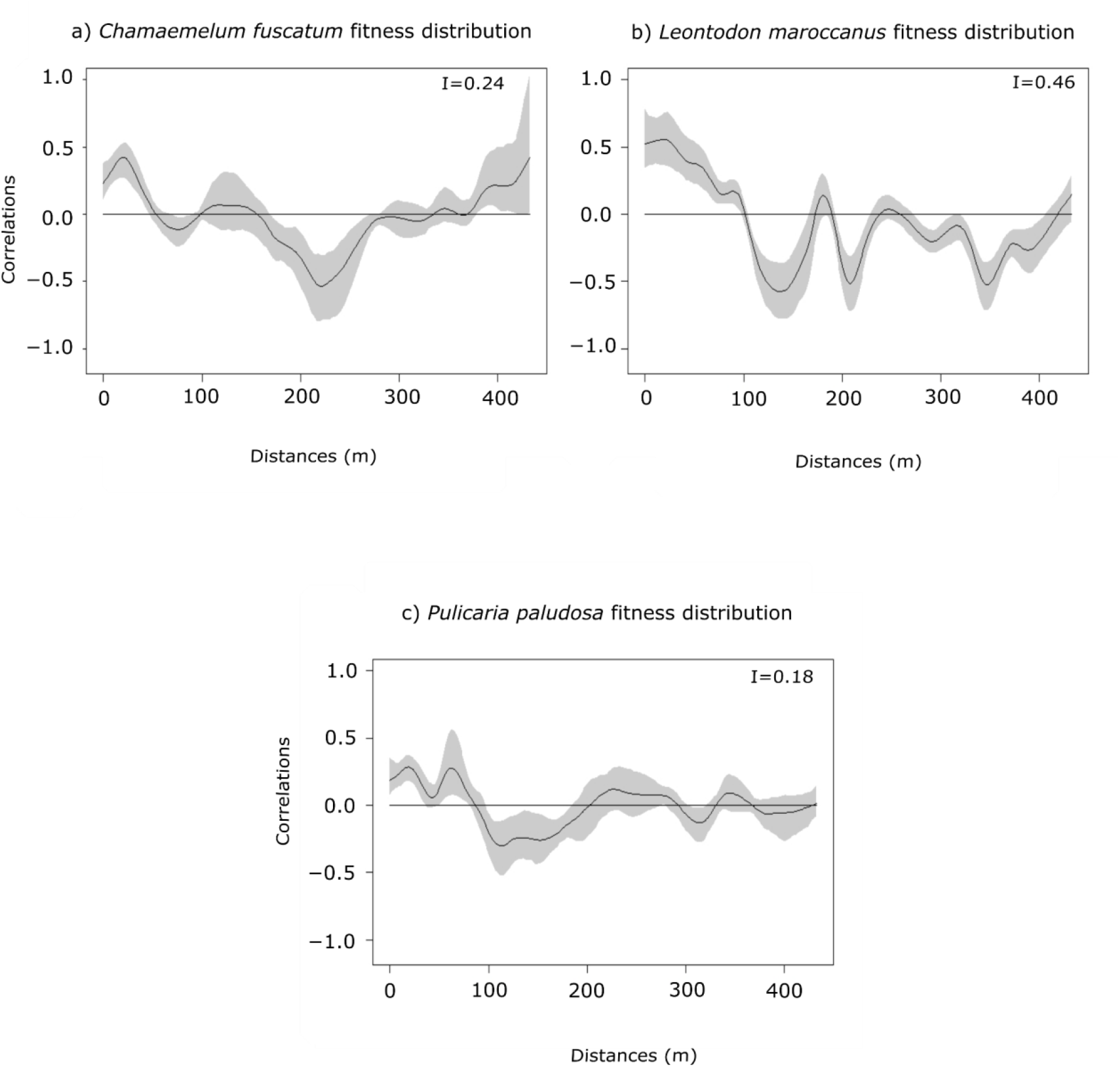
Spatial autocorrelation of fitness (reproductive success) distribution of plant species. The black line is the spatial correlation value that a species has for each distance, the grey shadow indicates the 95% of the confidence interval. The I values are the result of the Moran’s I statistic.

**Table A3.** This table shows the ANOVA result of each plant species with the constrained and the multigroup model. The “*” means that the result is significant, meaning that both models are not equal (if they are equal means that this species does not depend on the scale). We want to check if the models depend on the spatial scale (multigroup models). In the case of *C. fuscatum* and *L*.*maroccanus* the models that are more parsimonious (low AIC) are the multigroup and in the case of *P. paludosa* the most parsimonious model is the constrained.

| Chi-Squared Difference Test | DF | AIC | BIC | Pr(>Chisq) |
| --- | --- | --- | --- | --- |
| *C. fuscatum* multigroup | 48 | 5011.2 | 5399.0 | 1.58e-06* |
| *C. fuscatum* constrained | 108 | 5047.5 | 5219.9 |  |
| *L. maroccanus* multigroup | 44 | 5663.1 | 6117.7 | < 2.2e-16* |
| *L. maroccanus* constrained | 107 | 5790.8 | 5989.7 |  |
| *P. paludosa* multigroup | 32 | 2849.6 | 3289.2 | 0.45 |
| *P. paludosa* constrained | 95 | 2787.2 | 3003.5 |  |

The following equations specified in R are the models that we use to create the SEM for each species. Eq. (1), Eq. (2) and Eq. (3) are equal except for some particularities for each species. The “∼” sign means that there is a relation between the predictors, and the double sign “∼∼” means that there is a correlation between the variables, there is a covariation. It is important to remember that fruits in our study are the same as the number of flowers.

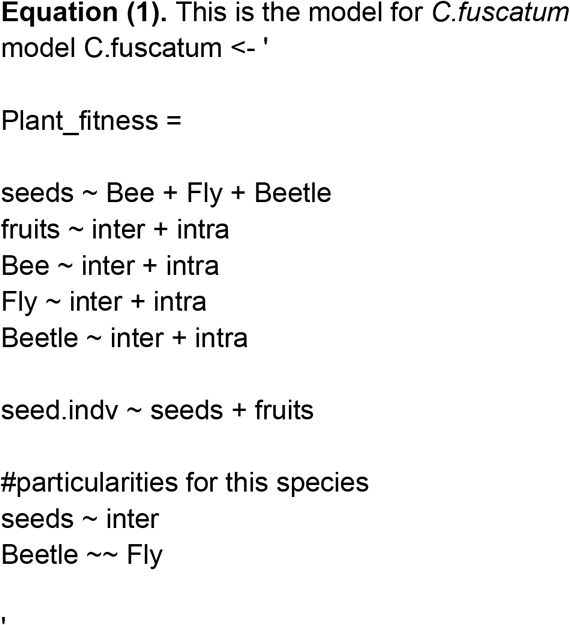

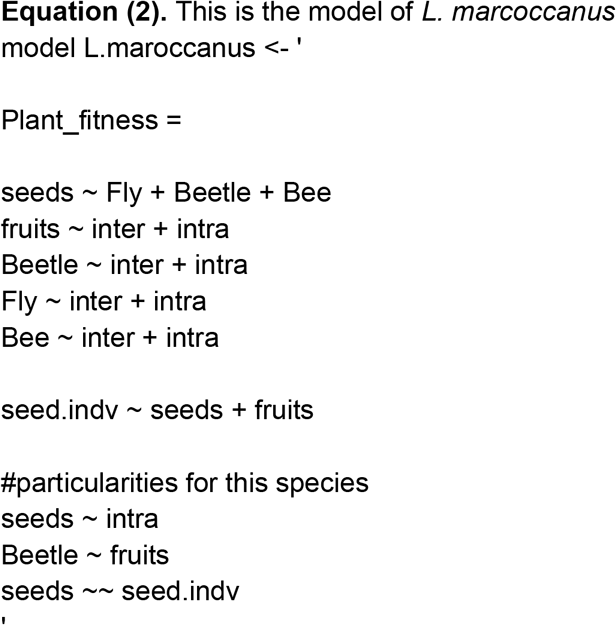

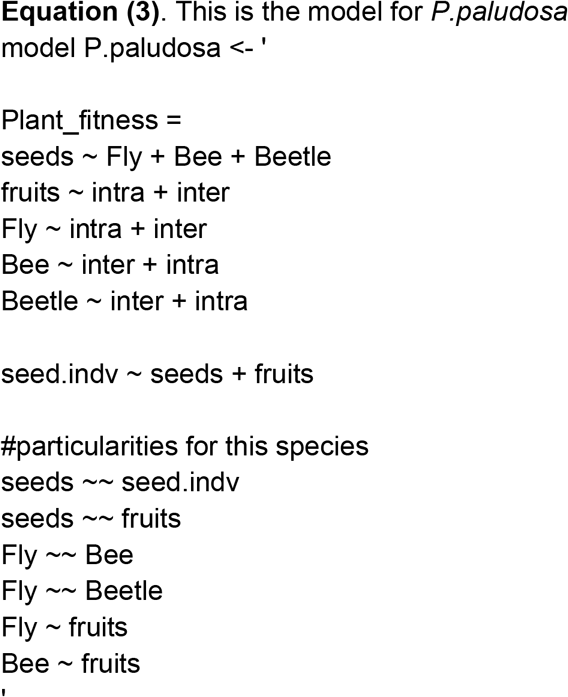

**Table A4.** Decomposition of the direct and indirect effects across the different scales in the species that are scale dependent (*C. fuscatum* and *L*.*maroccanus*). In the table it is shown the standardized total effects.

| Species | Scale | Total effect | Competition | Pollinators effect |
| --- | --- | --- | --- | --- |
| *C. fuscatum* | 7.5 cm | -0.217 | -0.227 | 0.010 |
| *C. fuscatum* | 1 m | -0.144 | -0.164 | 0.020 |
| *C. fuscatum* | 3 m | -0.352 | -0.370 | 0.018 |
| *C. fuscatum* | 6 m | -0.378 | -0.396 | 0.019 |
| *L. maroccanus* | 7.5 cm | -0.588 | -0.582 | -0.006 |
| *L. maroccanus* | 1 m | -0.070 | -0.071 | 0.001 |
| *L. maroccanus* | 3 m | -0.036 | -0.035 | -0.001 |
| *L. maroccanus* | 6 m | -0.041 | -0.040 | -0.001 |

